# Robust data-driven gene expression inference for RNA-seq using curated intergenic regions

**DOI:** 10.1101/2022.03.31.486555

**Authors:** Alessandro Brandulas Cammarata, Sara S. Fonseca Costa, Marta Rosikiewicz, Julien Roux, Julien Wollbrett, Frederic B. Bastian, Marc Robinson-Rechavi

## Abstract

RNA-Seq is a powerful technique to provide quantitative information on gene expression. While many applications focus on measuring expression levels, accurately distinguishing between actively and inactively transcribed genes is equally important for understanding gene function, development, and disease mechanisms. However, setting a biologically meaningful threshold for calling genes expressed is challenging due to variability in noise levels across different protocols, experiments or biological samples. We propose to define this threshold per sample relative to the background level observed in inactive genomic features, inferred by the amount of reads mapped to intergenic regions of the genome, and to call genes expressed if their level of expression is significantly higher than the estimated background noise. This approach can be applied to a single RNA-Seq library as well as to a combination of libraries from the same condition, in model and non-model organisms. We show that our method yields a more accurate prediction of expression state than existing methods, illustrated by consistent expression calls for biological replicates in the same tissue.

## Introduction

### Determining qualitative expression of a gene

Gene expression mediates biological function in two fundamental ways: by regulating the level of gene expression and by turning genes on or off in different conditions. While quantifying expression levels has been the primary focus of methodological developments^1^, accurately identifying whether genes are actively transcribed or inactive—the presence or absence of gene expression—is equally critical^2^. This distinction is essential for understanding gene function, developmental processes, disease mechanisms, and the intricate link between the genome and phenotype^3–9^.

The ability to determine which genes are active or inactive in specific conditions provides invaluable insights into gene regulation and cellular function. It enables researchers to annotate genes to specific conditions and facilitate integration across datasets or data types^10^. Moreover, prefiltering a dataset for genes that are present is an important first step in all transcriptome analysis workflows, including differential expression analyses (DEGs) and gene set enrichments^1,11^. Yet *ad-hoc c*riteria are often used although accurate presence/absence calls could reduce noise and enhance the power and accuracy of downstream analyses^12^.

However, the levels of expression of active genes can vary significantly between conditions, experiments, samples, as well as between model and non-model species (Figure 1). When comparing across species, these apparent differences are driven not only by biological divergence, but largely by disparities in genome annotation quality. For instance, non-model organisms often have many missing genes (especially non coding or short) or unannotated exons in otherwise annotated genes. This variability, which is caused by a mix of biological and technical factors, impacts the effectiveness of using a single fixed threshold on the expression levels, as routinely done in many studies^13,14^. This underscores the need for an adaptive biologically and statistically sound method to determine active gene expression.

**Figure 1:**
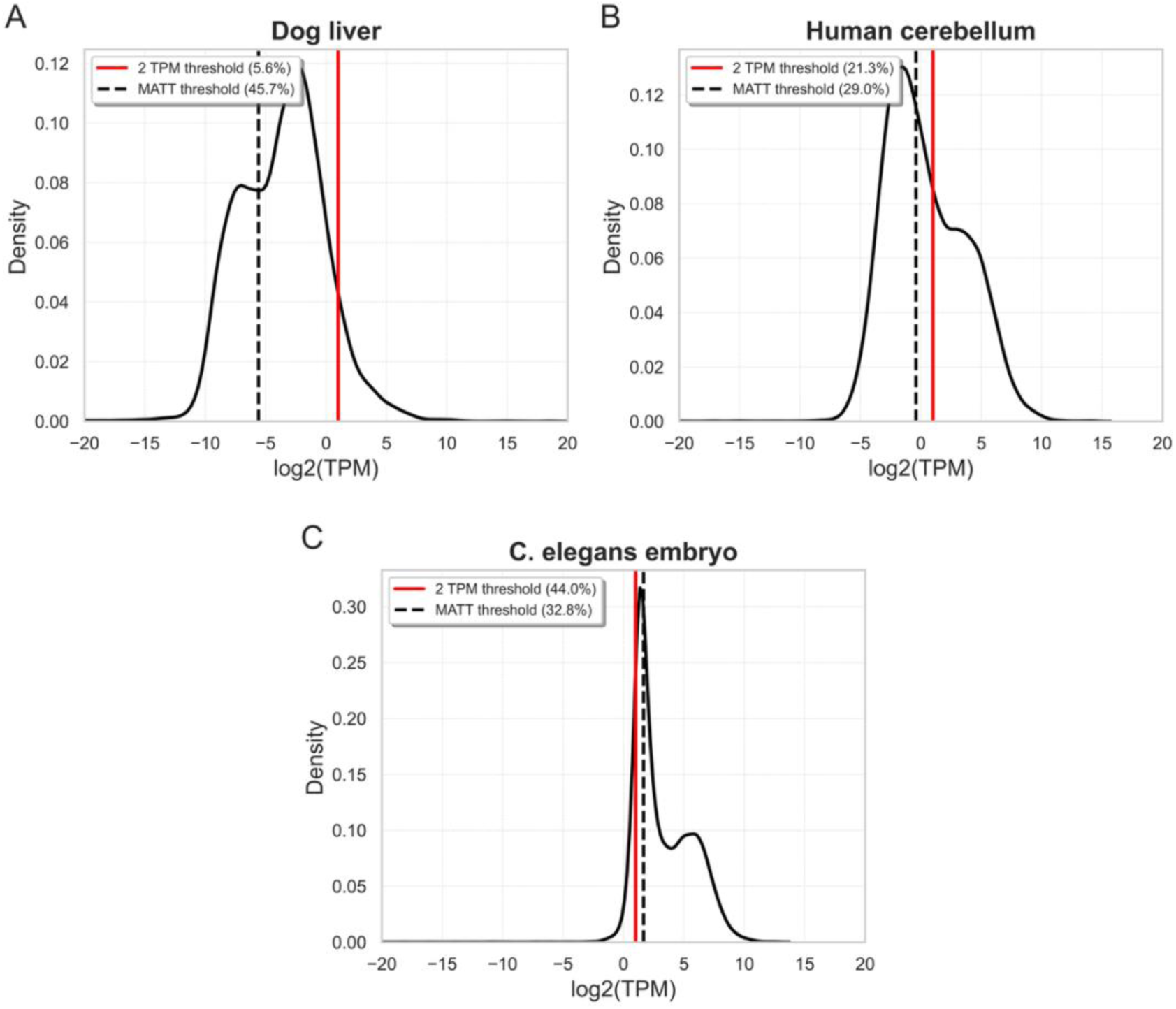
Density plots of log₂-transformed TPM values across annotated genes and expression thresholds in three RNA-Seq libraries. **(A)** dog liver (SRX8483838) **(B)** human cerebellum (SRX081989) **(C)** Caenorhabditis *elegans* embryo (SRX004863). The solid red vertical line represents the fixed threshold of 2 TPM commonly used in gene expression studies. The dashed vertical lines represent the library-specific MATT thresholds.

Transcription is inherently a noisy phenomenon^15–17^, influenced by active gene regulation, border effects from neighbouring active genes, stochastic binding of RNA polymerase^18–21^, and permissive chromatin states allowing leaky transcription^21–25^. Technical factors also introduce additional noise in gene expression measurements, such as sample contamination, RNA isolation methods and library amplification biases. Consequently, inactive genes can still produce RNA molecules through these various processes^26^, complicating the accurate identification of truly active genes.

### Current methods to call active expression

RNA-Seq is widely used as a powerful technology to quantify gene expression levels, using units normalizing for sequencing depth and gene length such as Transcripts Per Million (TPM), Reads Per Kilobase of transcript per Million mapped reads (RPKM)^27^, or Fragments Per Kilobase of transcript per Million mapped reads (FPKM). Arbitrary thresholds of these units are often employed in transcriptomics studies to call genes active or inactive, with little consensus on the exact value to use. Reported thresholds range from as low as 0.1 RPKM^28^ to as high as 3 TPM^29,30^. The Expression Atlas Baseline^31^ uses a threshold of 0.5 TPM or 0.5 FPKM to report expression. Many RNA-Seq studies even consider a gene expressed if any read is mapped to it^1,32^. There is no benchmark to evaluate the sensitivity and specificity at different thresholds.

Recognizing these challenges, researchers have explored methods to define sample-specific or gene-specific thresholds for distinguishing active and inactive genes. Hebenstreit et al.^26^ demonstrated that cultured murine Th2 cells exhibit two classes of genes—active and inactive—with some inactive genes receiving reads and contributing a clear left shoulder on the distribution of log-transformed RPKMs across genes. They identified gene classes by deconvoluting this distribution into two Gaussian components, validating gene status through RT-PCR and measurements of histone modifications. Wagner and Lynch^30^ proposed fitting a model to the raw TPM values, aiming to deconvolute a mixture of an exponential distribution for inactive genes and a negative binomial distribution for active genes. Hart et al.^33^ introduced an approach based on the distribution of RPKMs to model highly expressed genes as a Gaussian distribution and measure the distance to this distribution with a Z-score (zFPKM).

More recently, Thompson et al.^2^ proposed a Bayesian mixture model (implemented in the method “zigzag”) to infer active expression, formalizing the logic of Hebenstreit et al.^26^. Their model fits the log-transformed TPM values to a mixture of a Gaussian for inactive expression, one or more Gaussians for active expression (e.g., low and high expression), and a compartment of genes with no detected reads. They infer posterior probabilities for genes to belong to an inactive or an active component. To our knowledge, this represents one of the most advanced methods available to call genes as actively transcribed or not from RNA-Seq data. However, this approach presents practical limitations, such as the requirement of at least two libraries to perform the inferences, and computational convergence problems when there is high discrepancy in variance between samples (see Results). Other methods to call active expression have emerged in the last years, such as approaches leveraging Gaussian Mixture Modeling (GMM) such as GMMchi^34^ or BulkECexplorer^35^ but they cannot be used to get a consensus call across biological replicates and fail to produce accurate results at the single library level when distributions are not bimodal which is often the case in non-model species.

### Using non-genic regions of the genome to assess expression

All these methods rely solely on reads mapped to annotated genes. We propose that other genomic regions could also be informative, particularly for defining a background expectation for the class of inactive genes. Intergenic regions are genomic features that we do not expect to be actively transcribed, yet RNA-Seq libraries contain reads mapped to them due to technical and biological noise. Reads mapped to non-exonic regions, including intergenic regions, have already been shown to improve differential expression analysis^36^.

In this study, we introduce a novel approach that uses reads mapped to intergenic regions as a proxy for technical and biological noise, enabling us to establish sample-specific thresholds for detecting active gene expression in RNA-Seq data. Our method provides robust estimates of active gene expression across different experimental protocols and with both model and non-model species. It does not require replicates and is computationally efficient. By assessing the level of spurious intergenic expression —due to factors other than active transcription— we estimate the transcriptomic noise per sample and detect genes significantly actively expressed above it. This allows us to define a sample-specific false positive threshold for calling active expression.

By providing a more accurate and sample-specific approach to gene expression calling, our method enhances the reliability of downstream analyses and facilitates better integration across diverse datasets. Filtering for genes that are truly present improves the power and accuracy of transcriptome analysis workflows, including differential expression analyses. Our method is used for presence/absence calls in Bgee16^10^ and can be applied to any bulk RNA-Seq library through the BgeeCall Bioconductor R package^37^, making it readily accessible for widespread use in the research community.

## Results

### Robust background noise estimation

To accurately identify actively transcribed genes, we developed a method that uses reads originating from intergenic regions proximal to annotated genes. We use transcriptomic evidence to exclude putative unannotated genes or exons from our intergenic regions list (see Supplementary Material). By employing these filtered reference intergenic regions, we can estimate the background transcriptional noise specific to each RNA-Seq library. This allows us to define sample-specific thresholds—referred to as the Minimum Actively Transcribed Thresholds (MATTs)— for calling genes as actively expressed. Unlike fixed thresholds such as 2 TPM, which do not account for variability between samples, our method adapts to each library (Figure 1, supplementary figure 1). For example in a dog liver library, the MATT is lower than 0.1 TPM; in this case a fixed threshold of 2 TPM is too stringent and classifies only 5.6% of genes as active. Conversely, in a *C. elegans* library, the MATT is higher than the permissive threshold of 2 TPM, which includes 44% of genes.

### Robustness Across Diverse Experimental Protocols

To assess the robustness of our method across different experimental protocols and conditions, we processed RNA-Seq libraries coming from various human tissues and experimental labs, starting with the human whole blood samples obtained from the GTEx project^38^ curated in Bgee (full list of libraries used in supplementary files). Some of those libraries were prepared with hemoglobin depletion while others were not, but this information is not available in GTEx metadata, and thus it is not trivial to process them separately.

These blood samples showed a wide variation in the proportion of protein-coding genes called expressed with a fixed threshold —from 9.08% to 58.68%. In contrast, our method provided stable expression calls, consistently calling around 56% of protein-coding genes expressed across all blood libraries (44.77% to 67.78%; Figure 2A). This indicates that our method is robust to dramatic variations in biological and technical factors, for example sample handling procedures such as delayed handling or library preparation protocols. Indeed, the TPM distributions of those libraries show that the MATT adapts to the specific profile of each library.

**Figure 2:**
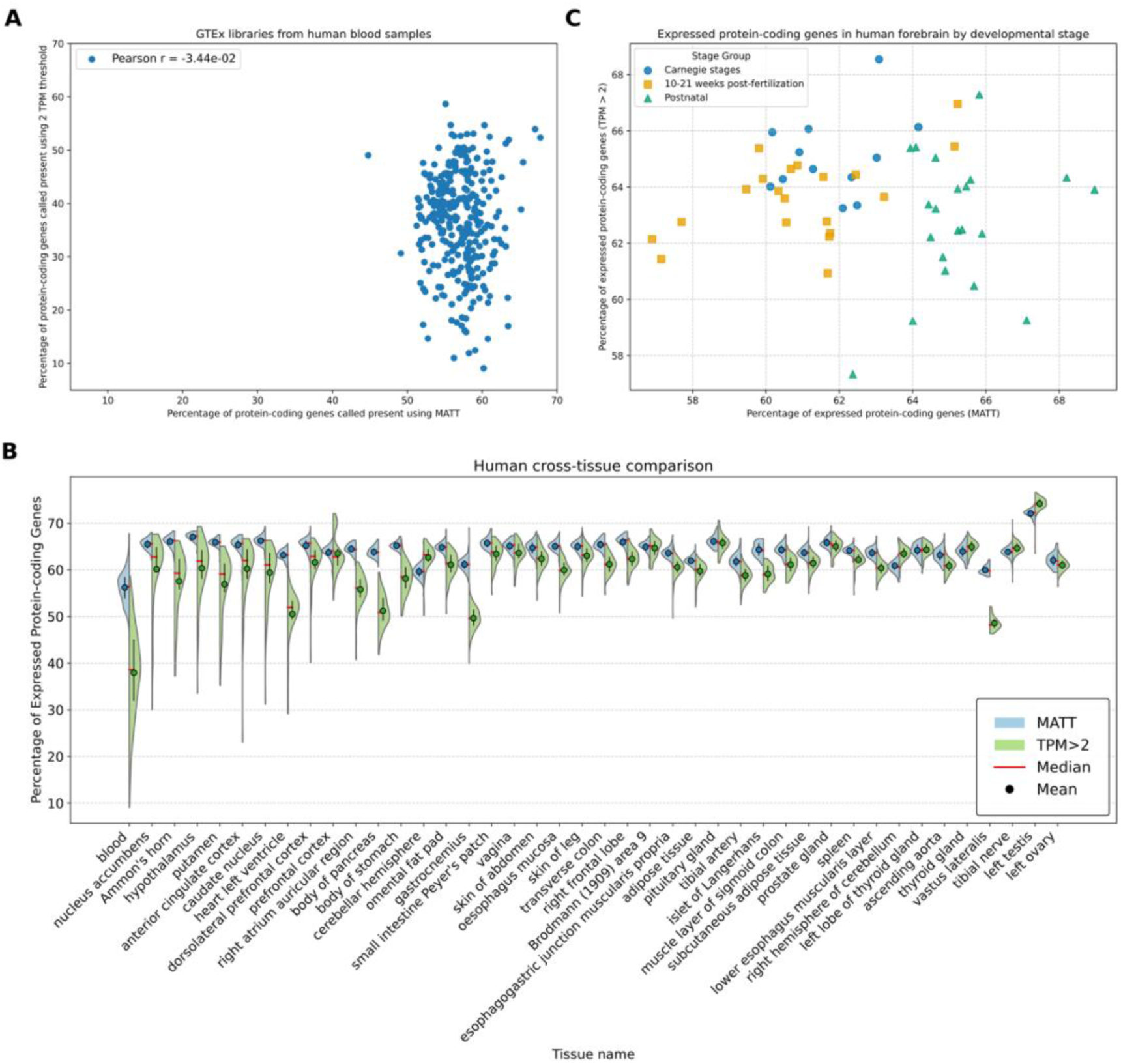
Comparison of proportions of protein-coding genes called expressed using MATT versus a fixed threshold of 2 TPM. **(A)** Comparison for human blood libraries. **(B)** Boxplots across different human tissues. Only tissues with more than 50 libraries are shown. For each tissue, the boxplots compare MATT (blue) and the fixed 2 TPM threshold (green). **(C)** Comparison across forebrain libraries. Blue and orange: embryonic developmental stages; green: post-embryonic developmental stages.

Next, we extended this analysis to multiple human tissues, from multiple experiments (including the whole GTEx version 6, see supplementary figure 2) to evaluate within-species variability between conditions across 7,400 RNAseq libraries. As seen for blood samples, using the MATT generally yields lower variance in the fraction of genes called expressed across libraries of the same tissue compared to using a fixed threshold, suggesting less susceptibility to technical variations and biological noise. A variance test confirms that the MATT results in significantly lower variability across libraries of the same condition compared to the fixed threshold method in 55 out of the 170 tissues (full F-test results in supplementary files). MATT shows higher variability in 11.

In the general case, we expect low variance of proportion of expressed genes between libraries from the same tissue, since they capture replicates of the same biology. But a fraction of the variance might be explained by remaining underlying biological variables. For example, in the forebrain, there is a high variance between libraries even with the MATT method, which is mostly explained by differences between embryonic (Carnegie stages and 10-21 weeks post-fertilization) and post-embryonic samples. Embryonic stages display a lower proportion of expressed genes than late stages with MATT, whereas no such pattern could be seen with the 2 TPM threshold method (Figure 2C).

### Robustness in Non-Model Species

To demonstrate the applicability of our method to non-model organisms, we analysed RNA-Seq data from 52 species and 30,056 RNA-seq libraries, from Bgee 16.0 (see supplementary tables 4 and 5 for the number of libraries and genome version used for each species). The pattern is similar to human tissues: MATT provides consistent expression call proportions within species, with reduced variance compared to the fixed 2 TPM threshold (Figure 3A). A variance test confirms that 25 species have lower variance with MATT, vs. only 6 with the 2 TPM threshold; for the 21 other species the test is not significant (full F-test results in supplementary files). The fixed threshold also shows more outliers. We also performed a cross-tissue variance analysis for pig, and as in the case of human data, MATT leads to more stable calls of expression within biological replicates (Figure 3B). Results are consistent comparing liver samples across 17 different species (Figure 3C).

**Figure 3:**
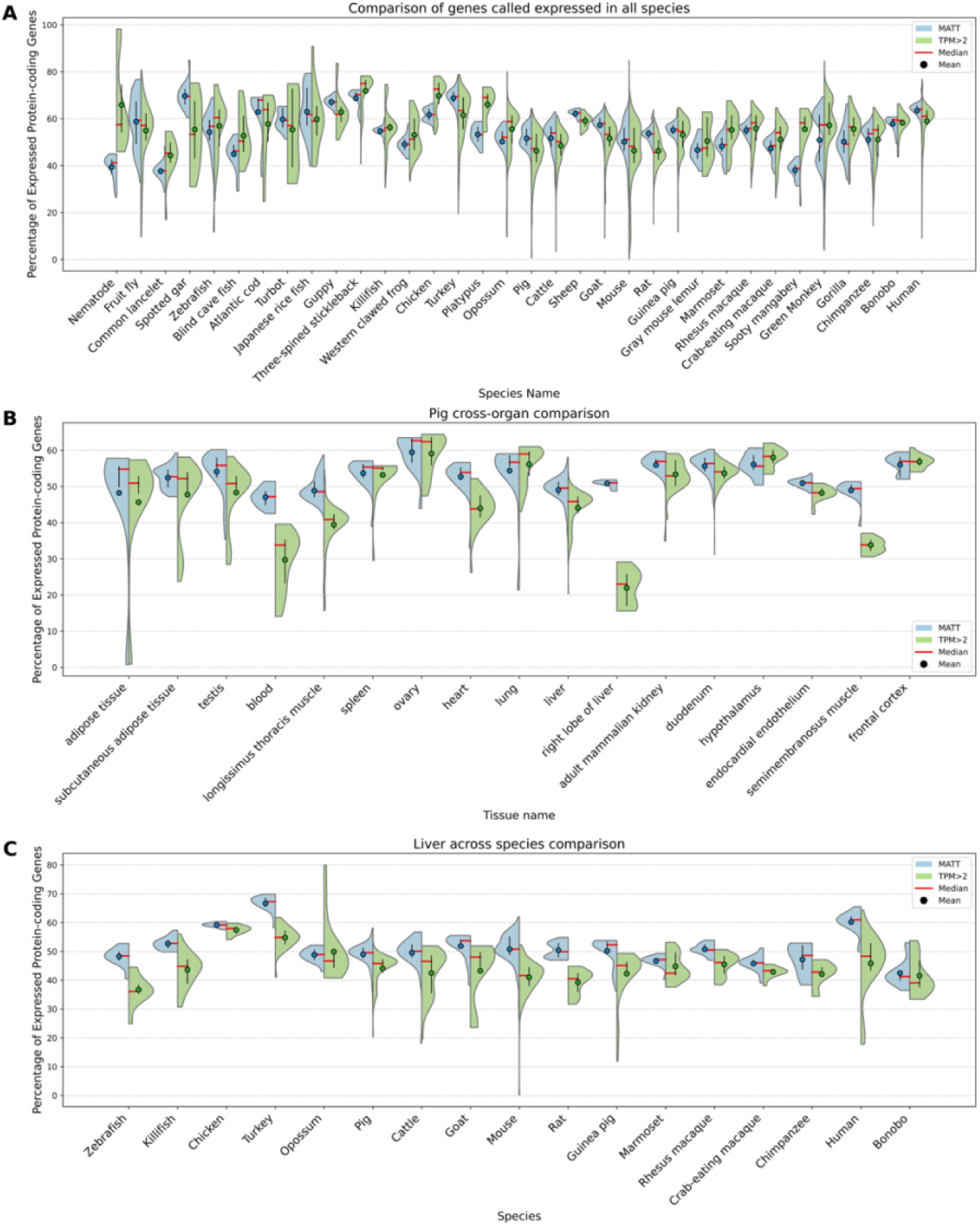
Performance of MATT and fixed threshold methods on non-model genomes. **(A)** Comparison of expressed gene proportions across different species and tissues using MATT and the fixed 2 TPM threshold. Only tissues with more than 15 libraries are shown. **(B)** Comparison across different pig tissues. Only tissues with more than 15 libraries are shown. **(C)** Comparison of liver across species. Only species with more than 15 libraries are shown. In all graphs, violin plots display the proportion of protein-coding genes called expressed using MATT (blue) and the fixed 2 TPM threshold (green).

Overall, our results on single library calls demonstrate that the MATT provides more accurate and consistent gene expression calls than the fixed 2 TPM threshold across various experimental conditions, tissues, and species.

### Calls over replicates

When multiple RNA-Seq libraries describe the same condition (biological replicates), either within one experiment or across experiments, it can be useful to make gene expression calls taking all the available libraries into account. i.e., to obtain one call per gene and per condition. For our MATT-based method, we combine the tests using the adjusted mean^39^ of library-specific p-values (see Supplementary Methods). Of note, we chose not to simply make calls per library, correcting for FDR over the libraries for a condition. Rather we combine the p-values, because we are not answering the question “is this gene expressed in at least one library?”, but “is this gene expressed in this condition, considering all of the available information?”. We compared these MATT-based calls to the Bayesian method zigzag^2^, which necessitates replicate libraries to make expression calls. We first evaluated method performance based on the two reference datasets used in the zigzag publication^2^, i.e. a set of genes inferred to be expressed in human lung based on 15 epigenetic markers (where expressed genes are defined as having active promoter and transcription mark but no repressed marks, and inactive genes as only repressed or heterochromatin marks), and a set inferred to be expressed in fly testis based on developmental genetic studies (specifically, 39 genes known to be active in the stem-cell niche of the testis against 119 assumed inactive odorant and gustatory receptors) (Figure 4A, B). Both methods were evaluated on the same dataset from Bgee libraries matching these conditions.

**Figure 4:**
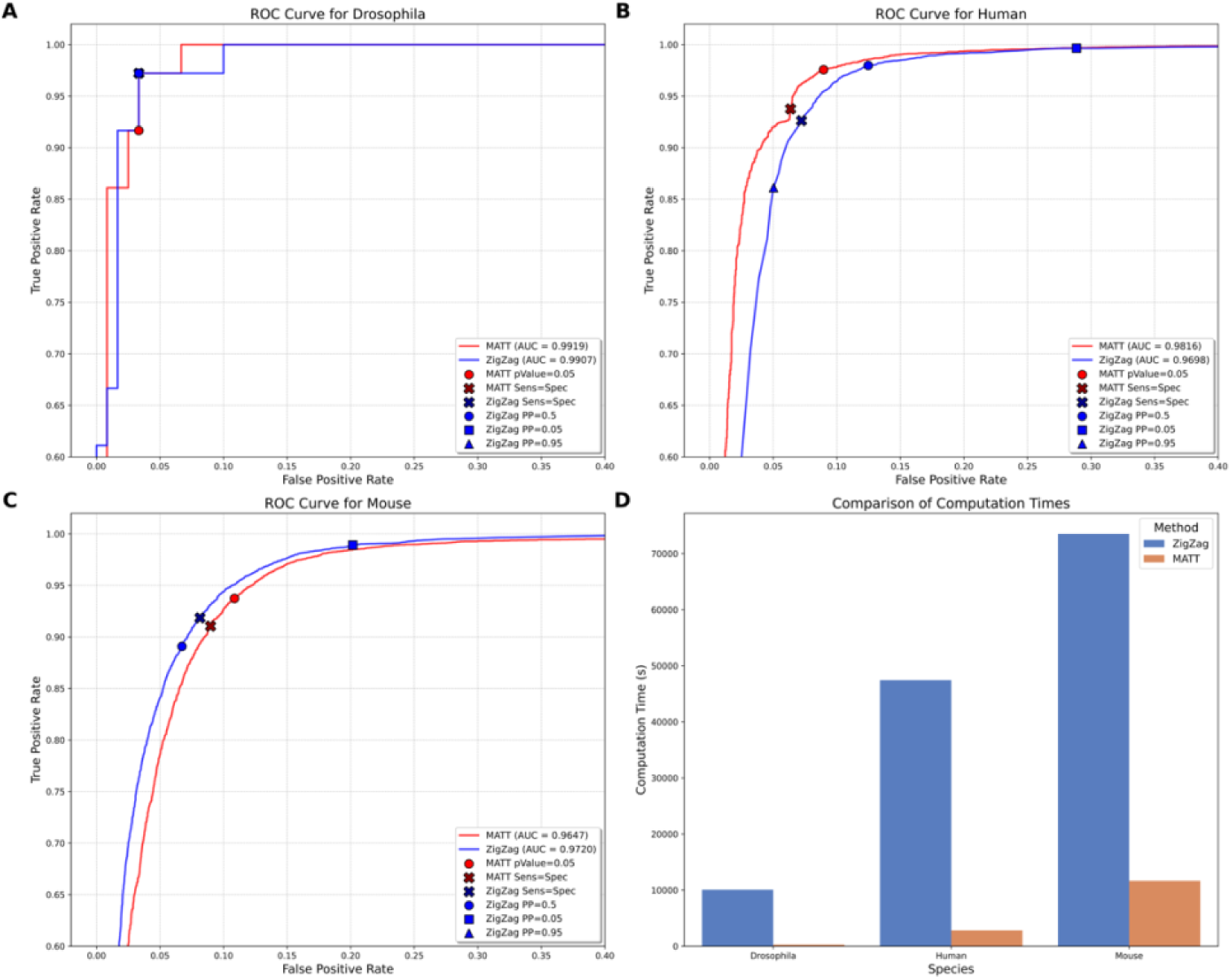
Comparison of the performance of MATT and zigzag to call active expression on a combination of libraries for 3 model species (*Drosophila melanogaster*, Human and Mouse). **(A)** Zoomed-in ROC curve of the calls of expression using 9 Drosophila testis libraries. **(B)** Zoomed-in ROC curve of the calls of expression for 67 human lung libraries. **(C)** Zoomed-in ROC curve of the calls of expression for 330 mouse liver libraries. The red lines show the performance of the MATT-based method and the blue lines show the performance of zigzag. For each method we estimated the inflection point (black cross), and we show the standard thresholds for each method as colored dots. **(D)** Comparison of computation times between MATT and zigzag for analysing RNA-Seq data from *Drosophila testis* (10 libraries), human lung (67 libraries) and mouse liver (232 libraries).

Both methods perform very well based on AUC, with the top scores for MATT (Figure 4A, B, supplementary figure 3A, B). For each method, we report two thresholds: the point where sensitivity equals specificity for inflection point calculated by using the coordinates of the TPR/FPR curve (supplementary tables 1, 2); and standard thresholds, i.e. for the Bayesian method posterior probabilities PP of 0.95, 0.5 and 0.05 (as shown in^2^) and for the MATT-based method a *p-value* of 0.05. With the inflection point thresholds, the false positive rates are low for all methods, with true positive rates of at least 92% (supplementary tables 1, 2). But these thresholds are not usable in practice, since we cannot know which threshold would maximize TPR/FPR for a new condition without external gene activity data. However, we observe that the thresholds of *p*=0.05 and PP=0.5, for MATT and zigzag, are close to those theoretical optimum inflection points. On the other hand, the widely used PP≥0.95 gives a very low TPR for zigzag.

We added an additional benchmark using mouse liver Ribo-Seq data. The rationale is that active translation by ribosomes provides a clear signal of activity of protein-coding genes that should only be seen in actively transcribed genes. Once again, both methods display very good AUC values (Figure 4C, supplementary figure 3C), and both performed very well using the inflection point threshold (supplementary table 3). At *p*=0.05 MATT has a bit higher FPR, while at PP=0.95 zigzag is again extremely conservative.

Even when their performances are similar, the MATT-based method is more computationally efficient (Figure 4D). Compared to zigzag, which relies on computationally intensive Bayesian inference and MCMC algorithms, MATT thus enables rapid analysis of large datasets. This increased speed facilitates its integration into existing workflows and makes it more practical for large-scale gene expression studies. Moreover, the MATT-based method can be used on a single library at a time (see previous sections), while zigzag requires replicates. Zigzag also failed to converge in some situations (see below).

We confirmed this pattern of similar accuracy but faster runtimes of the MATT-based method by subsampling 3 libraries per condition to infer expression calls (Figure 5). Of note, zigzag also failed to converge to a result for 2 Drosophila and 7 Mouse runs.

**Figure 5:**
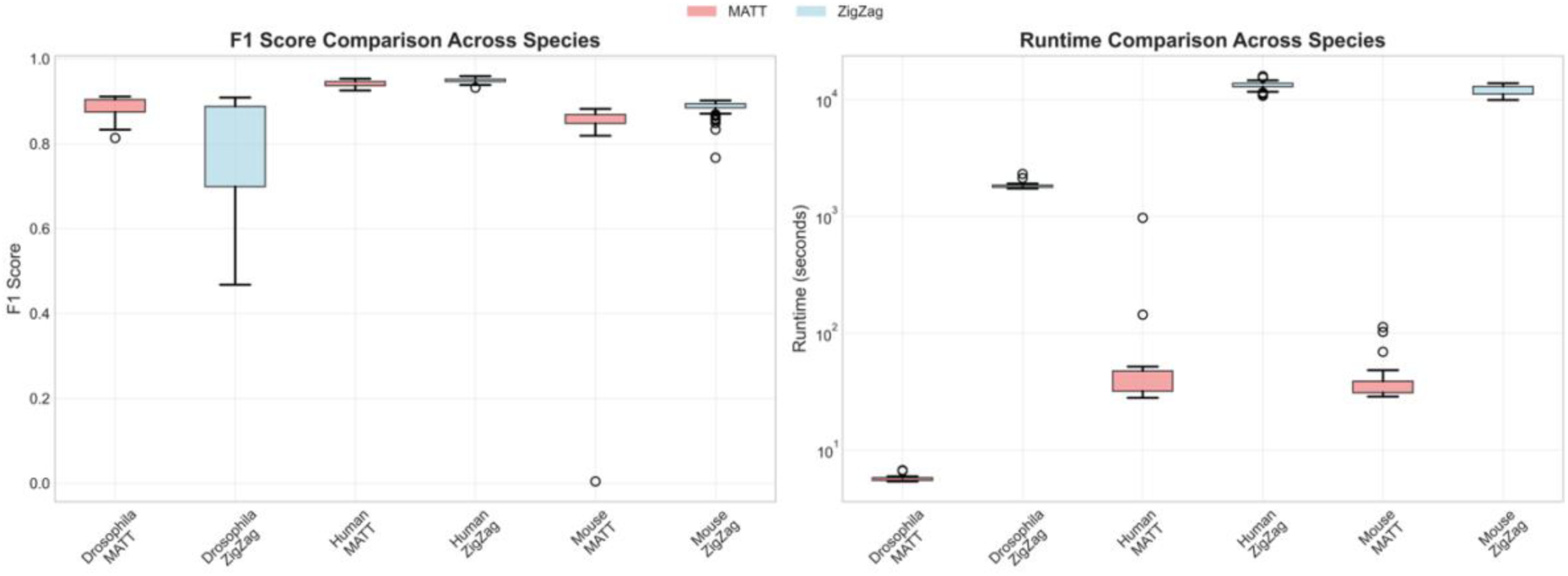
Comparison of the performance of the MATT-based method and zigzag to call active expression on a subsampling of 3 libraries per condition for 3 model species (Drosophila, Human and Mouse). **(A)** F1 score distributions. **(B)** Runtime distributions.

## Discussion

Determining when and where a gene is actively expressed is fundamental to understanding gene function, development, and diseases^4–9^. Accurately distinguishing between active and inactive genes in a given condition is crucial not only for biological interpretation but also for improving the reliability of quantitative analyses, such as differential expression studies^1,11^. Restricting analyses to expressed genes reduces noise from background transcriptional activity and enhances the power and accuracy of downstream analyses. However, inferring gene expression status from quantitative RNA-Seq data is challenging, as noted by Thompson et al.^2^: “Inferring the expression state (active or inactive) of a given gene from transcriptomic datasets is surprisingly difficult.” Ideally, a method to determine gene expression status should be robust to skewed expression distributions among genes, varying quality of genome annotations, and low numbers of replicates.

Several methods have been proposed that model the distribution of expression levels across genes to infer expression status^2,30,33,35,40^. In this work, we present an empirical approach that does not depend on such modelling but rather estimates the background transcriptional noise in RNA-Seq libraries by using reads assigned to intergenic regions. By defining a set of reference intergenic regions that are not actively transcribed, we can perform expression calls for individual genes in any sample, including those with only a single library.

We are not the first to compare RNA-Seq counts mapped between exonic and non-genic regions. Hebenstreit et al.^26^ observed that intronic regions exhibit intermediate expression levels between intergenic regions and exonic regions, consistent with the presence of immature or partially processed mRNAs and unannotated exons. While this observation is informative about transcriptional processes, it is less helpful for estimating background noise for expression calling. Hebenstreit et al.^26^ also used the 90th percentile of the intergenic TPM distribution as a threshold for expression calls. Since their study, the number of annotated non-coding RNA genes has increased substantially^41^, highlighting the importance of accurate genome annotations.

Subsequent studies have explored thresholds based on intergenic or intronic expression^42^, but these were limited in scope and did not provide general guidelines. Kharchenko et al.^43^ were among the first to use intergenic transcription to define gene expression thresholds, establishing a 0.4 RPKM cutoff to balance false positives and negatives in defining human housekeeping genes. However, the distribution of intergenic RPKM values was broad, again suggesting the inclusion of actively transcribed regions.

These findings, along with our own results, indicate that while intergenic regions can provide valuable estimates of background transcriptional noise, it is crucial to define reference intergenic sequences that are truly not actively expressed. This requires excluding unannotated genes and other potentially transcribed regions from the intergenic set. To define such reference intergenic regions, we leveraged the curated healthy wild-type libraries available in the Bgee database^10^. By excluding e.g. tumor samples or immortalized cell lines—where transcriptional regulation may be disrupted—we ensured that our reference intergenic regions represent genuine background expression. Including a diverse range of samples, representing different tissues and developmental stages, allows for a comprehensive assessment of intergenic expression across various conditions.

Our approach is readily extendable to other species. While we have primarily tested animal genomes, we anticipate that other large eukaryotic genomes with substantial intergenic regions would behave similarly. However, in compact genomes like those of yeasts or bacteria, the lack of extensive intergenic regions may limit the applicability of our method.

This consistency across both model and non-model species highlights the robustness of our method in accounting for library- and species-specific noise levels. By providing stable expression calls, our MATT method facilitates comparative studies and enhances the reliability of cross-species analyses.

One of the key advantages of our method is its robustness and computational efficiency. Unlike modelling methods^2^, which require multiple libraries and can be computationally intensive, our approach can be applied to individual libraries without the need for replicates. This makes it particularly suitable for studies with limited samples. Moreover, once the reference intergenic regions are defined for a species, they can be applied to any existing or new libraries from that species without additional recalibration. There is no need for posterior controls such as Markov Chain Monte Carlo convergence checks or resampling procedures, simplifying the workflow.

The sets of reference intergenic regions we compute may also have utility beyond expression calling. For example, they could serve as baseline references in epigenomic studies, such as providing controls for promoter histone mark analyses^44^. Furthermore, these regions could provide a valuable complement to existing Quality Control (QC) metrics, such as the RNA Integrity Number (RIN). By serving as an internal baseline for transcriptional noise or genomic DNA contamination, data from these intergenic regions could be integrated as features into machine learning models for automated sample QC and classification^45^.

Our method is also effective when combining information from multiple libraries. By aggregating p-values across libraries using methods like the mean adjusted by a factor of 2^39^, we can determine genes that are generally expressed in a particular condition. This is particularly useful when integrating data from different experiments, since active expression is more robust than level of expression^2^. Compared to a Bayesian approach, our method is more flexible, accommodating samples with high variance and different experimental protocols, and is more than 2 orders of magnitude faster.

In conclusion, we present a robust, versatile, and computationally efficient method, implemented as an open-source R package (BgeeCall), to distinguish actively expressed genes from background transcriptional noise in RNA-Seq data. By leveraging reference intergenic regions, our approach provides accurate expression calls that can be applied to single libraries or combined across multiple datasets, encompassing various experimental designs and protocols. By making this tool available, this method enhances the reliability of downstream analyses and facilitates better integration across diverse datasets, contributing valuable insights into gene regulation and function.

## Methods

### Data requirements

For each species under consideration, we initially require an annotated reference genome sequence and a set of RNA-Seq libraries. In practice, we used curated healthy wild-type libraries from the Bgee database^10^, but other sources can be used.

From the reference genome, we extracted two sets of genomic regions for read mapping: genic regions, which include all exonic sequences (both coding and non-coding), and putative intergenic regions. The definition of these putative intergenic regions depends on the quality of the genome annotation at this stage, i.e. they might include non-annotated genes or exons. To ensure that the intergenic regions have similar external noise factors as genes—such as chromatin compactness—we defined them as DNA regions located 0.5 kilobases (kb) away from the start or end of any protein-coding gene annotation and having a length of 1 kb (Figure 6A). These parameters can be adapted based on the structure and compactness of different genomes; in this study, we focused on animal genomes.

**Figure 6:**
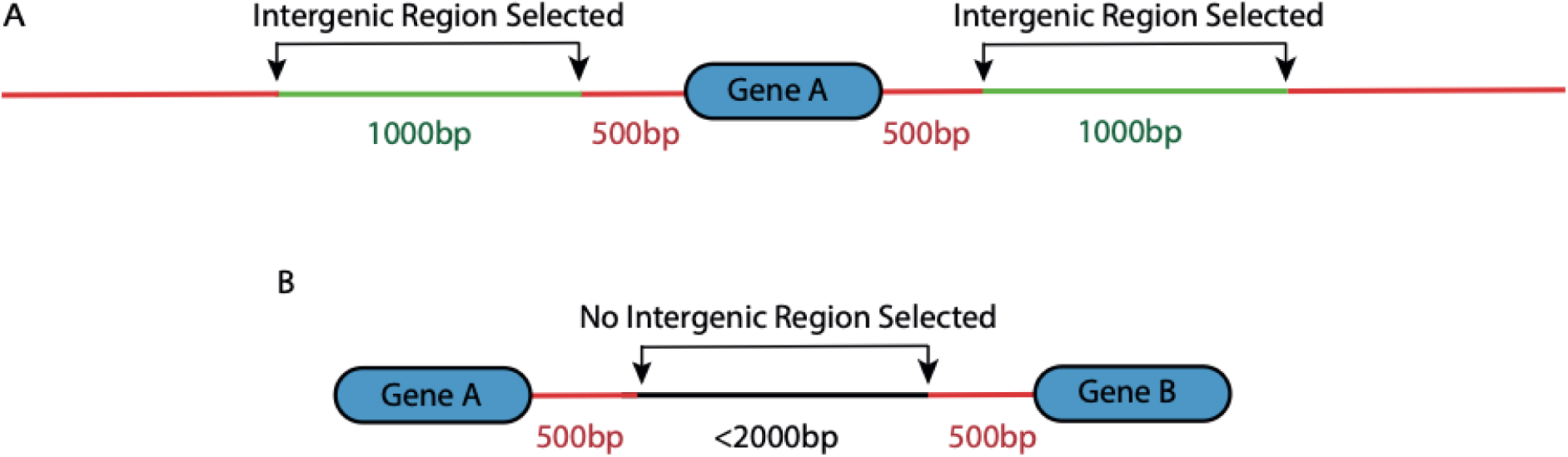
Definition of selected intergenic regions. **(A)** For each gene in the annotation, we select a 1 kb intergenic region upstream and downstream that is at least 0.5 kb away from the gene boundaries. Intergenic regions selected for analysis are shown in green; regions not included in the analysis are shown in red. **(B)** If two genes are too close to select an intergenic region for both (i.e., the required offsets overlap), no intergenic region is selected at that position.

In cases where two genes are too close to select an intergenic region for both, no intergenic region is selected at that position (Figure 6B). This approach prevents the inclusion of regions that may be strongly influenced by adjacent gene expression.

### Expression-based filtering of intergenic regions

Reads from each RNA-Seq library are then mapped to the genic and putative intergenic regions, to obtain the counts per region. We used Kallisto (version 0.46.0)^46^ to quantify transcript abundance from RNA-Seq data using pseudo-mapping, but the approach would be similar when using other read mapping tools such as Salmon^47^ or STAR^48^. For each gene, we summed the counts over all its transcripts to obtain gene-level counts.

Because of differences of genome annotation quality in the 52 species analyzed and to make sure that our intergenic regions are a robust estimator of non-actively transcribed regions, we filter the originally selected intergenic regions for regions which receive an unusually high amount of reads after summing counts from all available RNA-seq libraries (see supplementary methods “Intergenic region selection”).

### Sample-specific expression threshold

To detect actively expressed genes in individual RNA-Seq libraries, we perform a statistical approach based on a hurdle model with a one-sample one-tailed z-test to model non-zero values. This test determines whether a gene’s expression level is significantly higher than the background noise represented by intergenic regions. For each library, we define a sample-specific threshold, referred to as the Minimum Actively Transcribed Threshold (MATT_library_), which is used to classify genes as active or inactive and corresponds to a p-value of 0.05 in our one-sample test (see supplementary methods “Sample-specific expression threshold).

### Combining multiple libraries

For some applications, it is important to determine genes that are actively expressed across a set of RNA-Seq libraries. Therefore, when we have biological replicates we aggregate for each gene its significance p-value by averaging them across all replicates weighted by our confidence in the power of the library estimated by its sequencing depths (see supplementary methods “Combining multiple libraries”).

### Mouse liver benchmark

We use a processed mouse liver Ribo-Seq dataset downloaded from GEO (GSE67305) to benchmark methods both at individual library level and in the process of combining multiple libraries to call genes present and absent. Active translation by ribosomes provides a clear signal of activity of protein-coding genes, so true positives were defined as the union of protein-coding genes detected in all samples with ≥ 1 RPKM and of protein-coding genes detected in at least one sample with ≥ 5 RPKM. True negatives were defined as protein-coding genes which matched neither condition.

## Data access

All empirical data used in this work for RNA-Seq were obtained from Bgee 16 and are accessible through the Bgee website or BgeeDB R package^49^.

All input files and specific code to reproduce the figures or statistical files of this paper are available either in zenodo (https://zenodo.org/records/19347960) or GitHub (https://github.com/mrrlab/Robust_data_driven_gene_expression_inference_code). The full Bgee pipeline for RNA-Seq is available in GitHub at https://github.com/BgeeDB/Bgee_pipeline/tree/master/pipeline/RNA_Seq.

The BgeeCall R package used for methods in this work can be found in Bioconductor (https://bioconductor.org/packages/release/bioc/html/BgeeCall.html).

## Competing interests statement

The authors declare that they have no competing interests.

## Acknowledgments

We thank all the Robinson-Rechavi group for their useful discussions on the methods implemented on the paper. We acknowledge support by the SIB Swiss Institute of Bioinformatics, the Canton de Vaud, Swiss National Science Foundation grants 173048 and 207853, and the Swissuniversities Open Research Data program.

## Authors’ contributions

MR, JR and MRR designed the original approach. ABC, SSFC, JR, JW, FBB and MRR refined it. ABC performed the graphic visualization as well as tables for the paper. MRR wrote the first draft of the paper. ABC, FBB and MRR wrote the final version of the paper. All the authors contributed to result interpretation and discussion. All authors read and approved the final manuscript.

## Supplementary Material

### Sample-specific expression threshold

To detect actively expressed genes in individual RNA-Seq libraries, we perform a statistical approach based on a one-sample one-tailed z-test. This test determines whether a gene’s expression level is significantly higher than the background noise represented by intergenic regions. For each library, we define a sample-specific threshold, referred to as the Minimum Actively Transcribed Threshold (MATT), which is used to classify genes as active or inactive and corresponds to a p-value of 0.05 in our one-sample test.

We pseudoaligned reads to both transcripts and the reference intergenic regions, computing Transcripts Per Million (TPM) values per gene (by summing over transcripts) and per intergenic region, using Kallisto for quantification. We model the distribution of the log-transformed TPM values of the intergenic regions by fitting a normal distribution.

We first identify intergenic regions that are outliers to ensure a robust estimation of the background expression distribution. Specifically, we calculate the interquartile range (IQR) of the log₂(TPM) values of the intergenic regions and remove any intergenic regions with log₂(TPM) values more than 1.5 times the IQR above the third quartile or below the first quartile.

After outlier removal, we calculate the mean (μ) and standard deviation (σ) of the log₂(TPM) values of the remaining intergenic regions, assuming that these values follow a normal distribution representing the background noise.

For each gene i in the library, we compute a z-score to quantify how many standard deviations its expression level is above the mean of the intergenic regions:

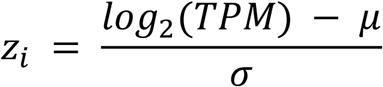

We then calculate a p-value for each gene based on the null hypothesis that the gene’s expression level originates from the same distribution as the intergenic regions (i.e., the gene is not actively expressed). Since we are interested in whether the gene’s expression is significantly higher than the background, we perform a one-tailed test:

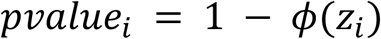

where *ϕ*(*z*_*i*_) is the cumulative distribution function of the standard normal distribution evaluated at *z*_*i*_. This p-value defines the first part of the hurdle model that evaluates the probability for intergenic regions to have an expression higher than a desired TPM value. The second part simply models the probability for intergenic regions to not receive zero reads and is equal to the ratio of intergenic regions having at least 1 read assigned to them. To get the final p-value of the hurdle model, we multiply the p-values of both parts.

The library-specific TPM threshold to call genes expressed, referred to as the Minimum Actively Transcribed Threshold (*MATT*_*library*_), is defined as the smallest TPM value where the p-value is less than or equal to a chosen significance level α (we use α = 0.05 unless otherwise specified):

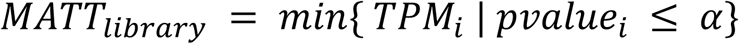

Genes with TPM values greater than or equal to *MATT*_*library*_ are considered actively expressed in that library. This approach allows us to classify genes as active or inactive based on a statistical assessment relative to the background noise, tailored to each individual library.

### Combining multiple libraries

For some applications, it is important to determine genes that are actively expressed across a set of RNA-Seq libraries. To combine information from multiple libraries, we tested two different methods (Table 1). Our approach involves combining the library-specific p-values using the mean of the values adjusted by a factor of 2, following the method described by Vovk and Wang^39^. While the adjusted mean of p-values is used in this paper, other methods, such as multiple test correction, are available in the BgeeCall R package and should be used when the question is about whether a gene is expressed in at least one sample instead of being generally expressed in this condition.

**Table 1:** Two methods are compared in this study: our method, implemented in the BgeeCall R package, which calls expressed genes at the individual sample level and then combines information from multiple libraries; and the zigzag method from the literature, which combines libraries using a Bayesian inference approach. Our approach uses a reference set of intergenic regions to call expressed genes.

| Method | Calls per library | Approach used to combine libraries | Reference | Availability |
| --- | --- | --- | --- | --- |
| BgeeCall | Yes | adjusted arithmetic mean | this work | BgeeCall package |
| zigzag | No | Bayesian inference | Thompson et al.<sup>2</sup> | zigzag package |

We compared the performance of our method with the Bayesian method zigzag developed by Thompson et al.^2^ (R package version: zigzag_0.1.0; repository: https://github.com/ammonthompson/zigzag). The zigzag method inherently requires multiple libraries to infer gene expression states and was tested across different conditions such as developmental stages, sexes, or strains.

To validate these approaches and calculate true and false positive rates, we used the same datasets as Thompson et al.^2^, specifically one for human lungs that contains active/inactive genes defined from epigenomic data. Another from fly testis containing a gold standard set established from genetic evidence. Additionally, we included a ribosome profiling dataset from Mus musculus (mouse) liver^50^ (see method section for details).

In our p-value approach, we combine information from multiple libraries by applying the following merging function M{1, K} to the p-values, where K denotes the number of libraries analysed for a given condition:

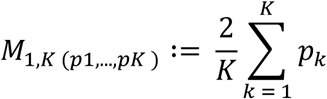

This function adjusts the combined p-value to account for the number of libraries, as described by Vovk and Wang^39^. Genes are then classified as actively expressed across the set of libraries if the combined p-value meets the significance threshold (e.g., p ≤ 0.05).

We also implemented the option in the BgeeCall package to weight the different libraries p-values. In those cases, the p-values are calculated by using the weighted arithmetic mean, with either the sequencing depths or, as implemented in Bgee, the number of unique TPM values as the weight of each library. However, as described by Vovk and Wang, if one of the library weights is bigger than half the sum of all weights then the merging function will be the following:

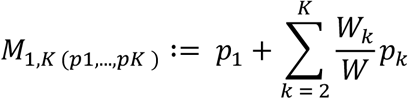

### Intergenic region selection

Publicly available genomes often contain ambiguous bases coded as ‘N’, which can impact read mapping and downstream abundance quantification. For the 52 species present in Bgee 16, the proportion of ‘N’s varies from 0% in Caenorhabditis elegans to 35.5% in Gadus morhua. These ‘N’s are frequently found in blocks of up to thousands of bases and predominantly affect intergenic regions. Since the absence of genome annotation is the only criterion to generate candidate intergenic regions, reads will map poorly to regions containing ‘N’s, potentially biasing abundance estimates. For instance, Kallisto pseudo randomly transforms all ‘N’ bases to A, T, C, or G bases during the pseudo-mapping step.

To mitigate this issue, we generated intergenic regions of fixed length (1 kb) located at least 0.5 kb away from the start or end of any gene annotation (Figure 6A). We excluded any regions that contained ambiguous bases ‘N’, ensuring that all selected intergenic regions are free of such bases. This filtering step is crucial to avoid biases in read mapping and quantification due to ambiguous sequences. The resulting set is available at [https://www.bgee.org/ftp/intergenic/2.1/]. We recommend performing similar filtering for any new genome to be analysed.

While some well-annotated species (e.g., human, mouse, fruit fly) have only a very small fraction of intergenic regions with high expression levels, other species, often non-model organisms, present large fractions of intergenic regions with high expression levels (supplementary figure 4), in general species with a higher amount of non protein-coding genes annotated have less intergenic regions excluded. This is likely due to the poor annotation of transcribed elements such as non-coding genes, very short coding genes or unannotated exons.

To define a strict set of intergenic regions, excluding these actively expressed regions, we compared the distributions of expression levels (log₂(TPM)) between intergenic regions and protein-coding genes. We calculated kernel density estimates (KDEs) for both distributions using the aggregated TPM values across all libraries for a given species.

For each TPM value, we computed the probability density from the KDEs for both the intergenic regions (P_intergenic_(TPM)) and the protein-coding genes (P_genic_(TPM)). We then identified the TPM value at which the probability density of the intergenic regions becomes consistently lower than that of the protein-coding genes. Formally, this threshold is the smallest TPM value above which P_intergenic_(TPM) < P_genic_(TPM) for all higher TPM values. This point represents the threshold above which intergenic regions are more likely to be actively transcribed, possibly due to unannotated genes or other transcribed elements. We define this TPM value as the species-specific maximum inactively transcribed threshold, or MITT_species_ (see supplementary files for per species filtering of intergenic regions).

The final set of reference intergenic regions consists of all intergenic regions with TPM values below MITT_species_ (see supplementary table 6 for the number of intergenic regions excluded per species). These regions are considered to represent the background level of technical and biological noise in RNA-Seq data for the species. Intergenic regions with TPM values above MITT_species_ are excluded from further analyses, as they may represent unannotated genes or regions of active transcription.

## Supplementary Tables

**Supplementary table 1:** Benchmark of combining multiple libraries for human lung data. Abbreviations: True positive (TP), False positive (FP), True negative (TN), False negative (FN), True positive rate (TPR), False positive rate (FPR).

| Method | Threshold | Number samples | Precision | TP | FP | TN | FN | TPR | FPR |
| --- | --- | --- | --- | --- | --- | --- | --- | --- | --- |
| BgeeCall | inflection point<br>($p \leq 0.0304$) | 67 | 0.9601 | 6767 | 281 | 4138 | 450 | 0.9376 | 0.0636 |
| zigzag | inflection point<br>($PP \geq 0.8710$) | 67 | 0.9544 | 6685 | 319 | 4100 | 532 | 0.9263 | 0.1002 |
| BgeeCall | $p \leq 0.05$ | 67 | 0.9468 | 7041 | 396 | 4023 | 176 | 0.9756 | 0.0896 |
| zigzag | $PP \geq 0.5$ | 67 | 0.9277 | 7071 | 551 | 3868 | 146 | 0.9798 | 0.1247 |

**Supplementary table 2:**
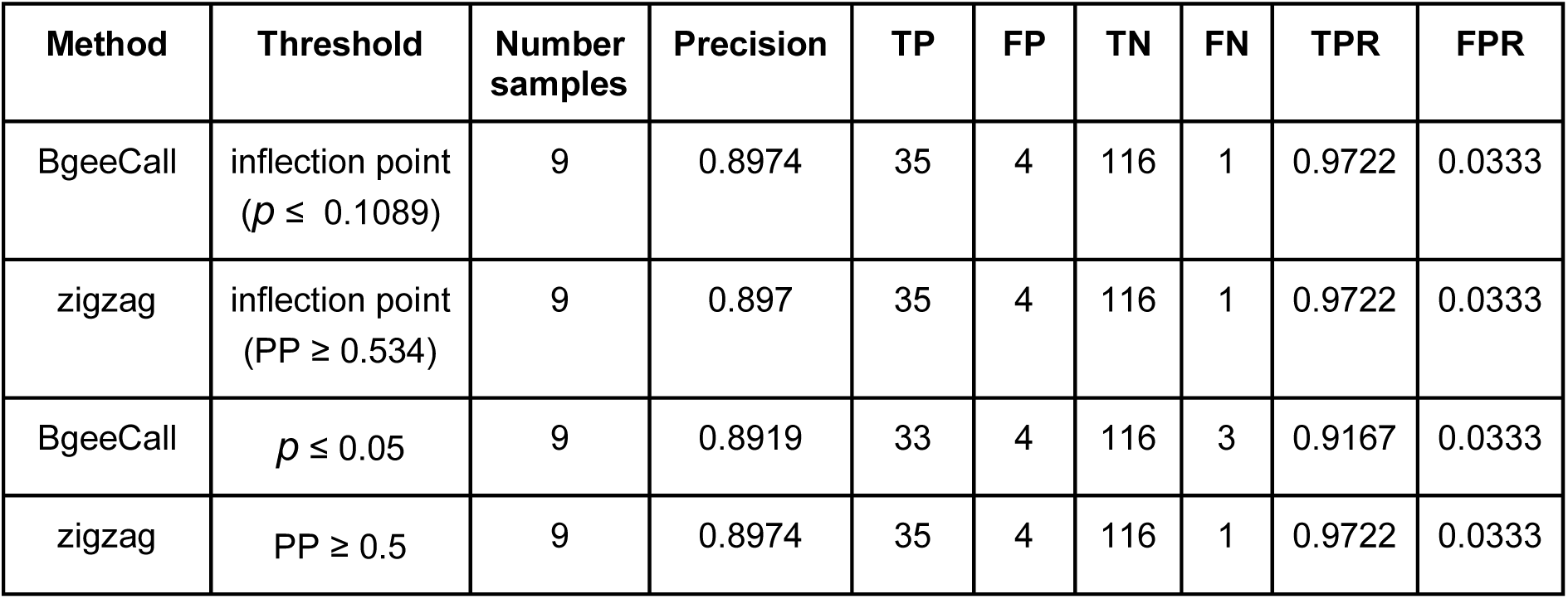
Benchmark of combining multiple libraries for fly testis data. Abbreviations: True positive (TP), False positive (FP), True negative (TN), False negative (FN), True positive rate (TPR), False positive rate (FPR).

**Supplementary table 3:**
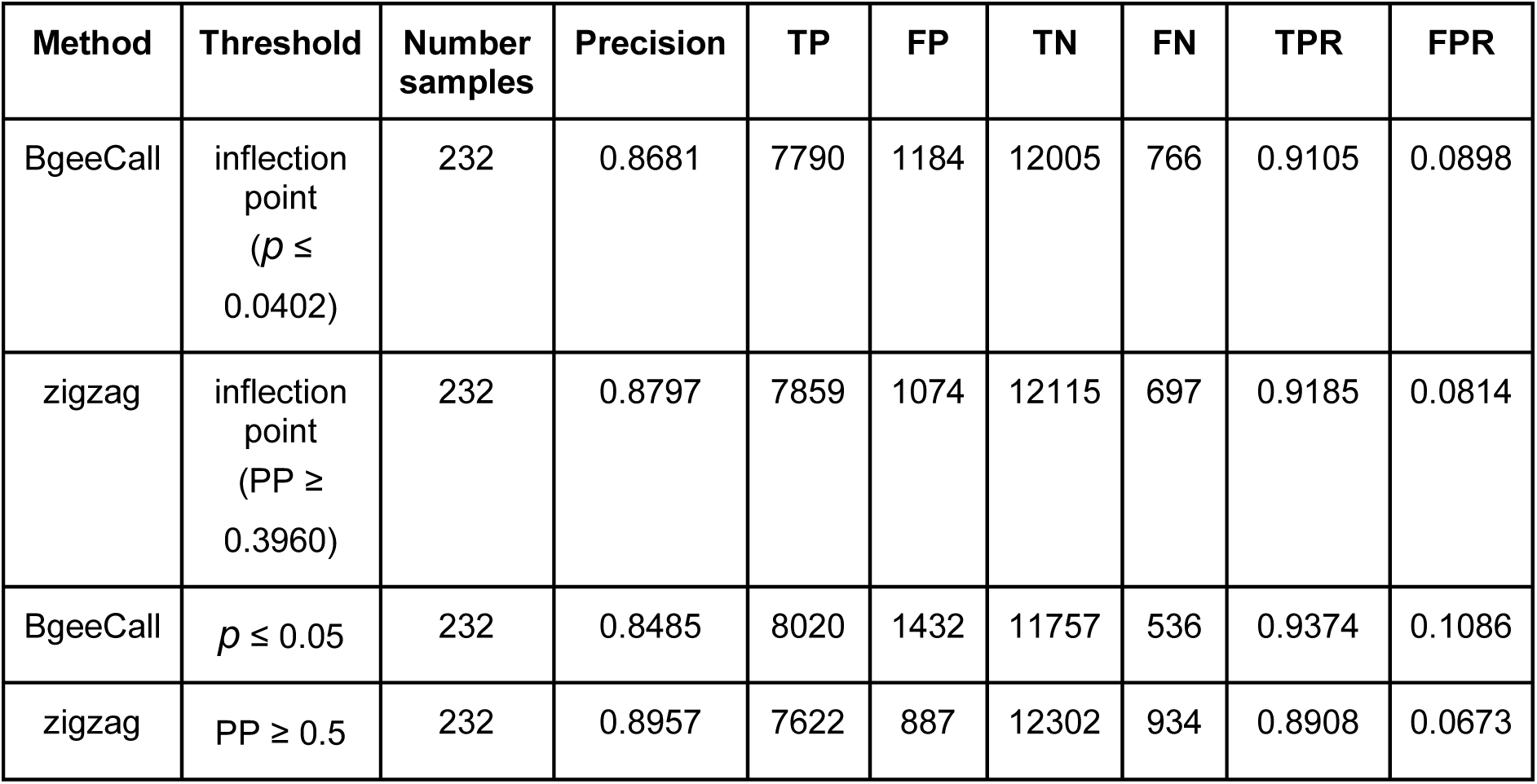
Benchmark of combining multiple libraries for mouse liver data. Abbreviations: True positive (TP), False positive (FP), True negative (TN), False negative (FN), True positive rate (TPR), False positive rate (FPR).

| Method | Threshold | Number samples | Precision | TP | FP | TN | FN | TPR | FPR |
| --- | --- | --- | --- | --- | --- | --- | --- | --- | --- |
| BgeeCall | inflection point<br>($p \leq 0.0402$) | 232 | 0.8681 | 7790 | 1184 | 12005 | 766 | 0.9105 | 0.0898 |
| zigzag | inflection point<br>(PP $\geq 0.3960$) | 232 | 0.8797 | 7859 | 1074 | 12115 | 697 | 0.9185 | 0.0814 |
| BgeeCall | $p \leq 0.05$ | 232 | 0.8485 | 8020 | 1432 | 11757 | 536 | 0.9374 | 0.1086 |
| zigzag | PP $\geq 0.5$ | 232 | 0.8957 | 7622 | 887 | 12302 | 934 | 0.8908 | 0.0673 |

**Supplementary table 4:**
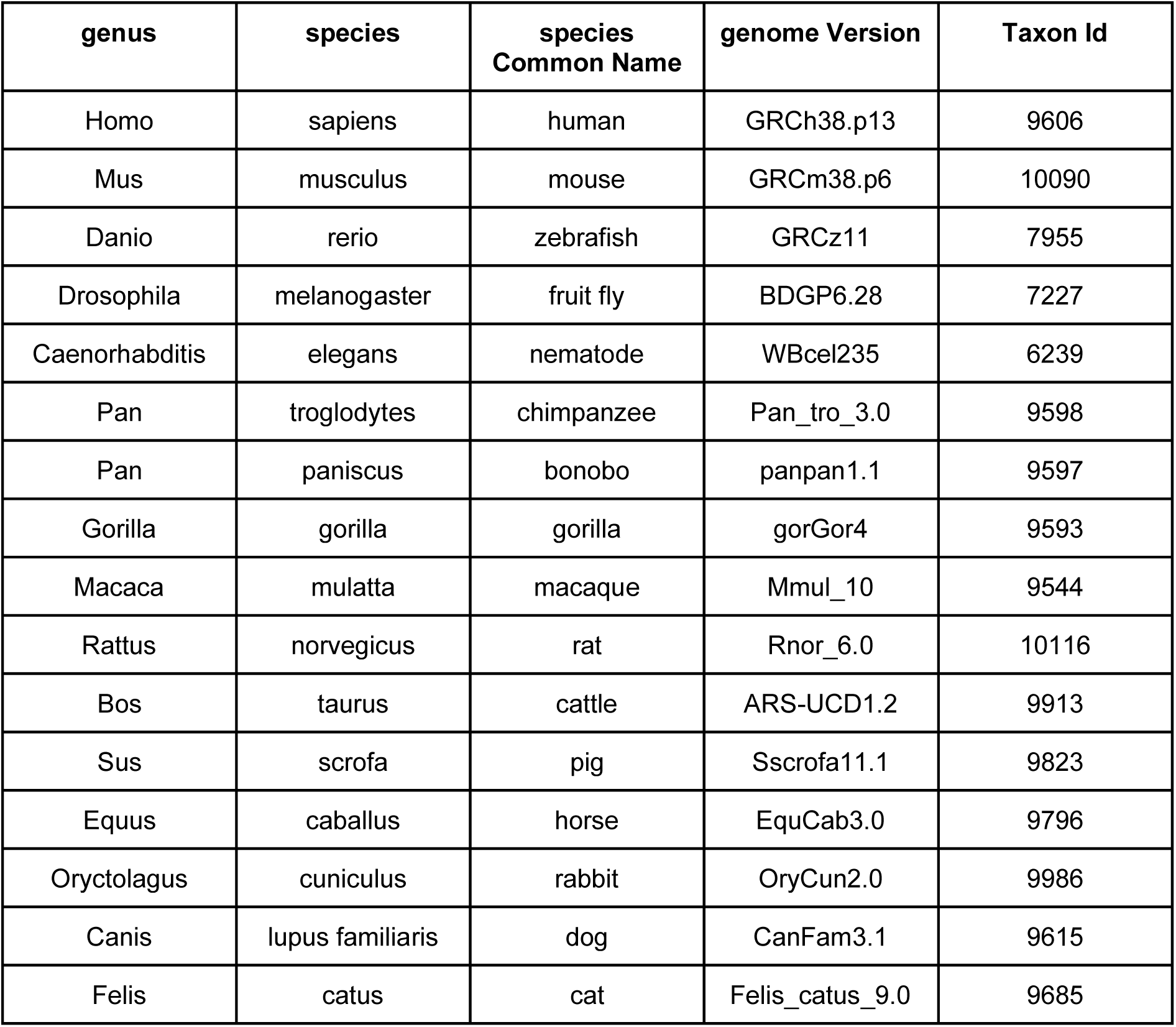
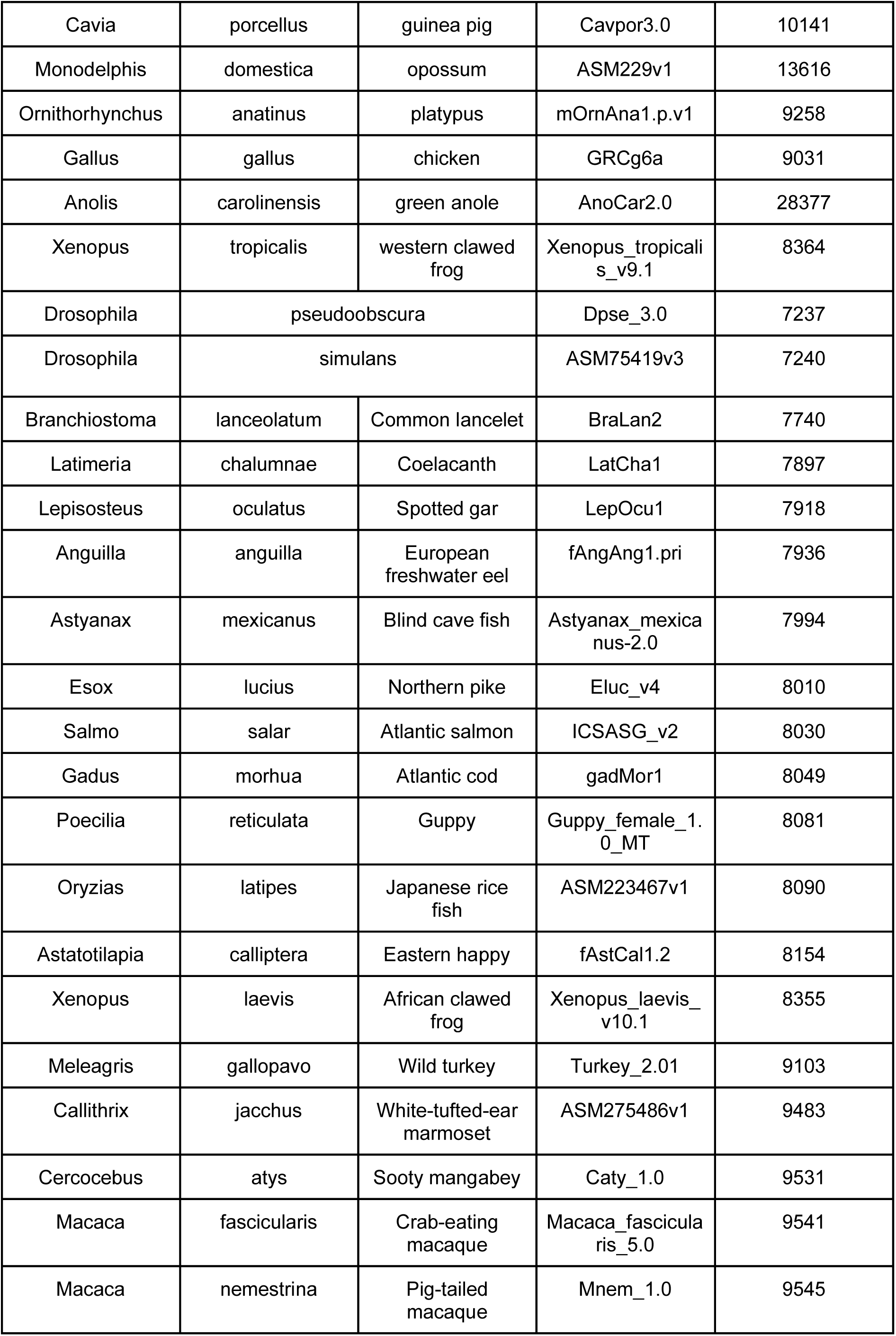
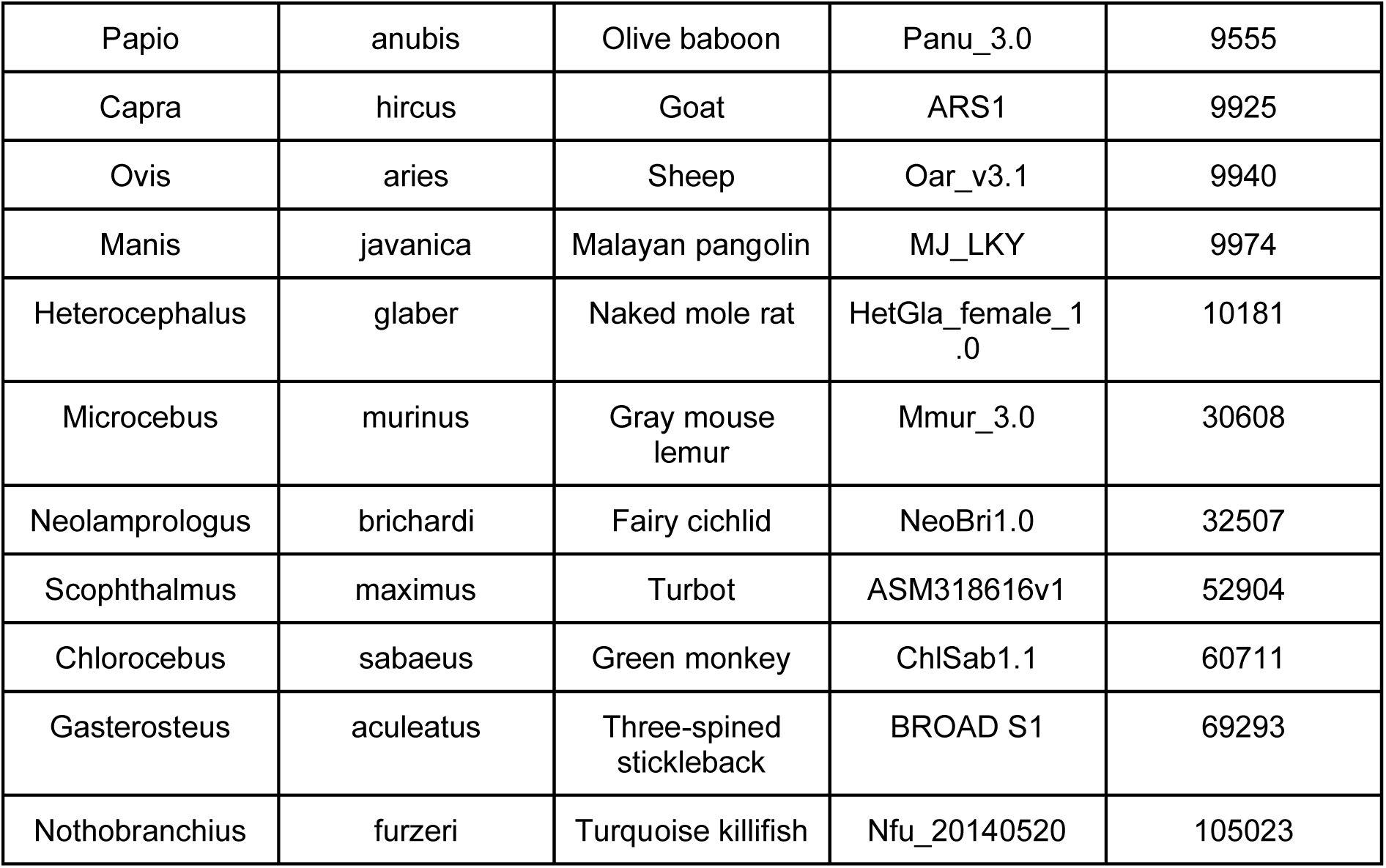
Genome versions used for results in the paper taken from ensembl^51^ release 113 and ensembl metazoa release 60.

**Supplementary table 5:**
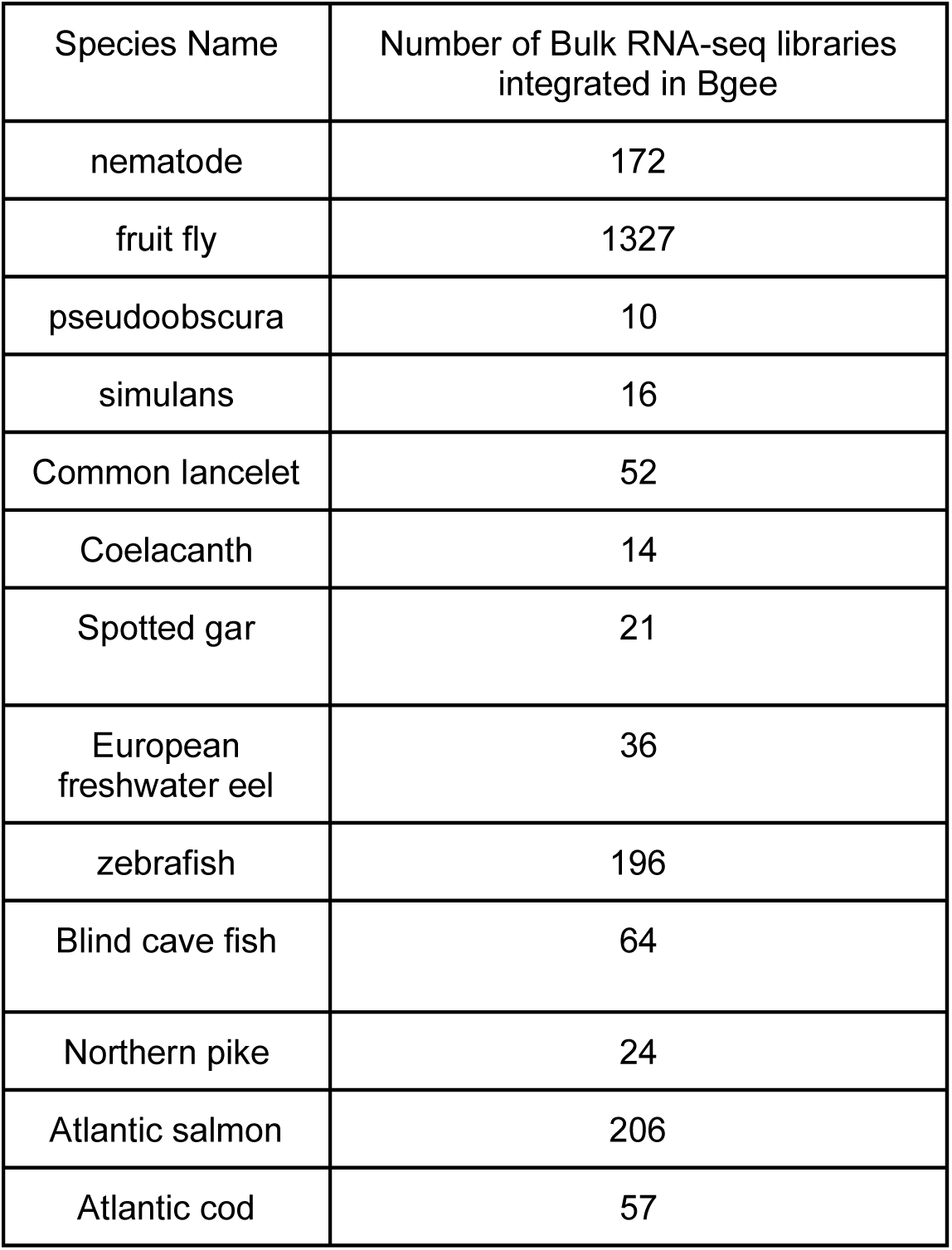
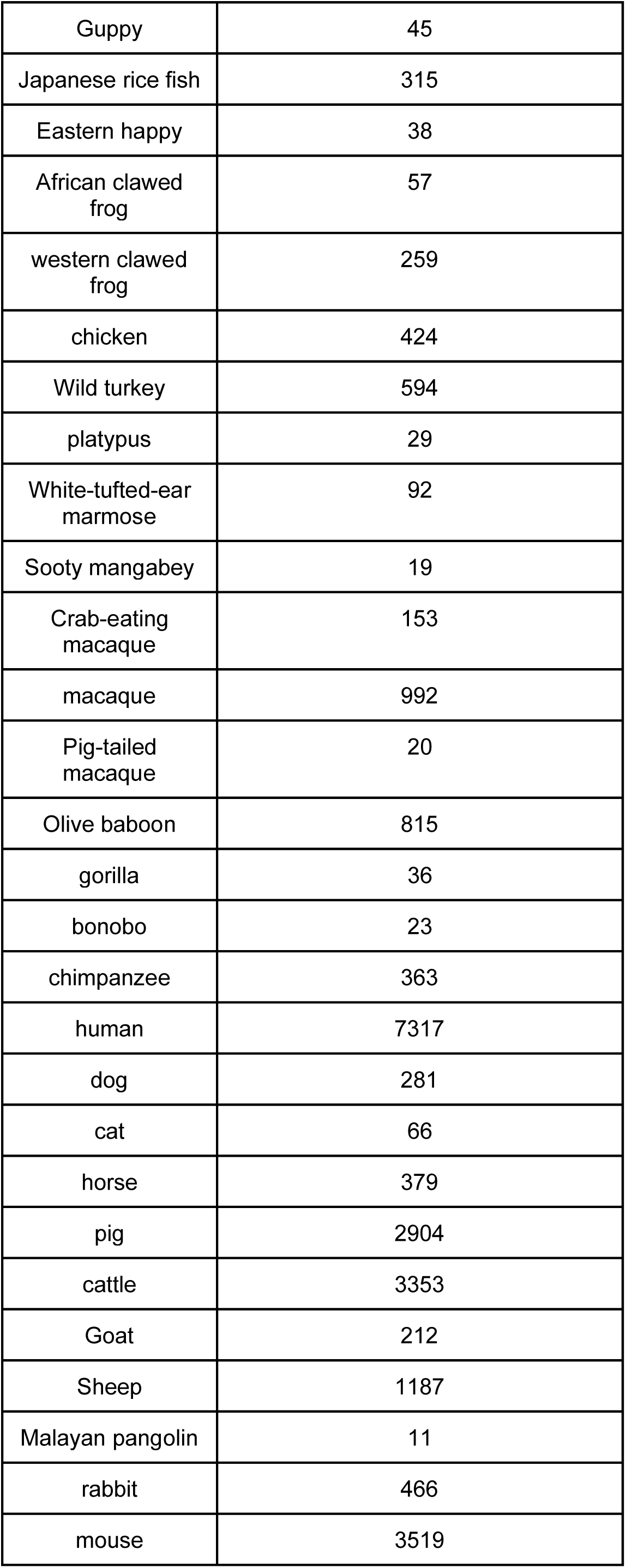
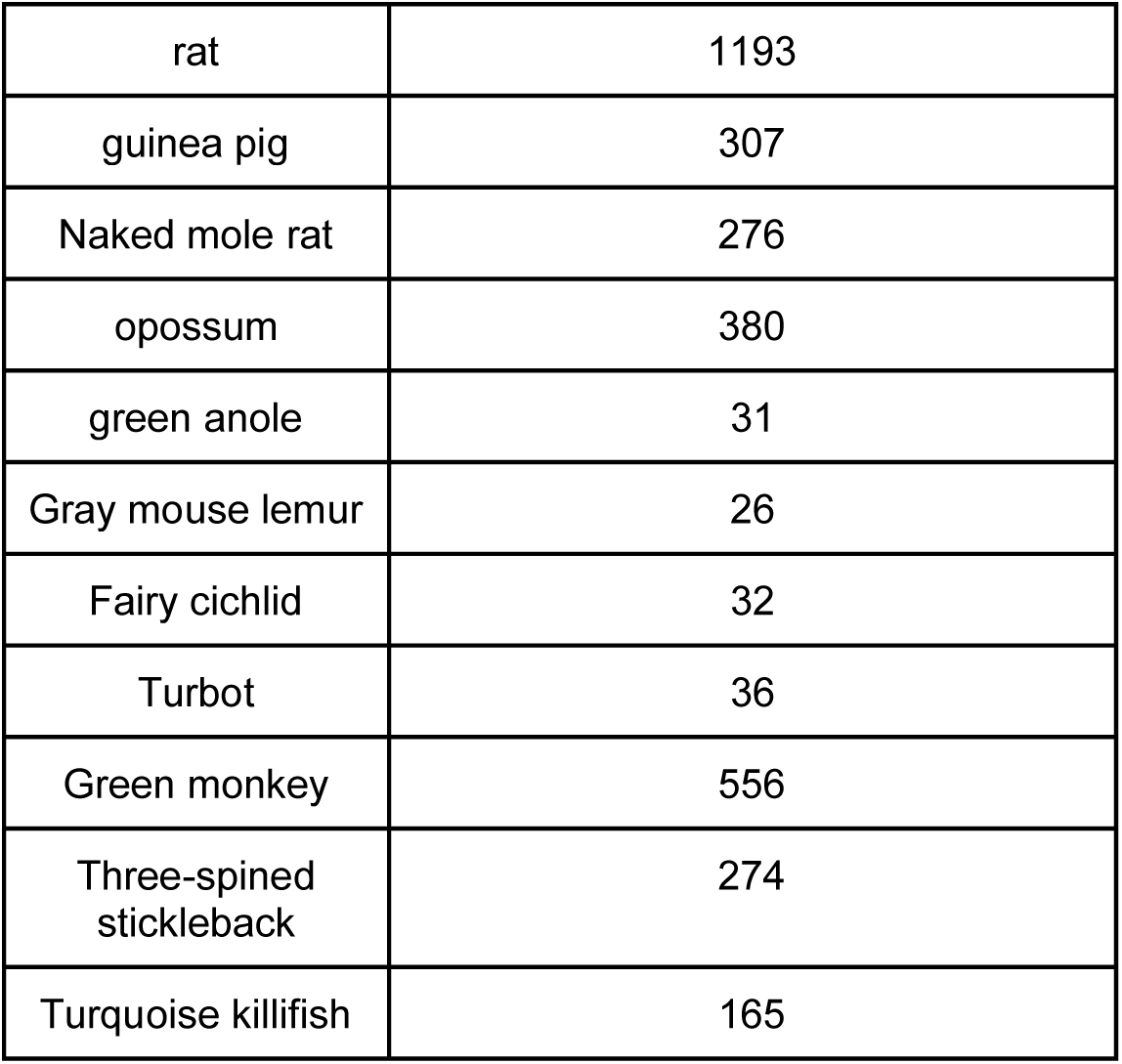
Number of RNA-Seq libraries annotated and integrated in Bgee 15 for all 52 species.

**Supplementary table 6:** Number of protein-coding, non-coding, reference intergenic and non-reference intergenic regions per species in Bgee 16 release.

| Species Name | Number of reference intergenic regions | Number of excluded intergenic regions | Number of protein-coding genes | Number of non-coding genes |
| --- | --- | --- | --- | --- |
| Sheep | 23682 | 5148 | 20921 | 6133 |
| Turquoise killifish | 18060 | 4938 | 23421 | 2054 |
| Crab-eating macaque | 21985 | 3041 | 22323 | 16227 |
| Blind cave fish | 24041 | 8063 | 26662 | 758 |
| rat | 27577 | 2945 | 23098 | 7464 |
| Naked mole rat | 23535 | 3859 | 23189 | 12139 |
| Eastern happy | 20939 | 4820 | 25713 | 7628 |
| Goat | 26751 | 5148 | 21343 | 5928 |
| White-tufted-ear marmose | 23305 | 4926 | 21965 | 18223 |
| fruit fly | 3299 | 65 | 13986 | 10292 |
| chicken | 10381 | 1691 | 17007 | 13101 |
| dog | 21866 | 2067 | 20500 | 10621 |
| cattle | 25103 | 3022 | 23739 | 12336 |
| Wild turkey | 13893 | 3983 | 16226 | 1744 |
| Atlantic cod | 15826 | 5105 | 23510 | 5553 |
| Turbot | 13361 | 4604 | 21260 | 13193 |
| Coelacanth | 19195 | 3814 | 19569 | 3059 |
| Green monkey | 25641 | 4425 | 19165 | 8820 |
| gorilla | 25089 | 3640 | 21587 | 8497 |
| Drosophila simulans | 4090 | 446 | 14179 | 1367 |
| macaque | 23573 | 3990 | 21591 | 13841 |
| human | 13580 | 248 | 20116 | 58816 |
| Drosophila pseudoobscura | 4347 | 651 | 14574 | 2722 |
| rabbit | 23142 | 3726 | 20599 | 8988 |
| Guppy | 16676 | 6103 | 22857 | 397 |
| mouse | 18504 | 914 | 21748 | 56550 |
| guinea pig | 21135 | 3617 | 18095 | 8760 |
| platypus | 21608 | 3184 | 17417 | 8855 |
| horse | 19563 | 1553 | 21426 | 17423 |
| European freshwater eel | 17830 | 5136 | 25903 | 5548 |
| Pig-tailed macaque | 23777 | 3367 | 20872 | 7993 |
| Malayan pangolin | 21512 | 1774 | 20586 | 12432 |
| Gray mouse lemur | 21855 | 3378 | 18816 | 7982 |
| Fairy cichlid | 15149 | 6493 | 23568 | 843 |
| zebrafish | 22137 | 4758 | 25432 | 7088 |
| Spotted gar | 15482 | 5461 | 18341 | 4974 |
| chimpanzee | 27728 | 3788 | 23302 | 10427 |
| pig | 26634 | 1208 | 22018 | 13664 |
| nematode | 2378 | 209 | 19985 | 26941 |
| African clawed frog | 40749 | 5482 | 34476 | 9980 |
| Japanese rice fish | 18508 | 4349 | 23600 | 765 |
| bonobo | 24369 | 3395 | 21041 | 9217 |
| Atlantic salmon | 39067 | 3958 | 47043 | 22346 |
| Sooty mangabey | 24282 | 3115 | 20746 | 7713 |
| Common lancelet | 22629 | 8890 | 35955 | 0 |
| western clawed frog | 22372 | 3179 | 22102 | 2512 |
| opossum | 25582 | 5039 | 21380 | 13605 |
| cat | 23036 | 4095 | 19564 | 9986 |
| Olive baboon | 23563 | 5423 | 21731 | 12104 |
| Three-spined stickleback | 9840 | 2410 | 22341 | 8075 |
| green anole | 22034 | 5272 | 21554 | 4925 |
| Northern pike | 21017 | 4379 | 25514 | 5450 |

## Supplementary Figures

**Supplementary figure 1:**
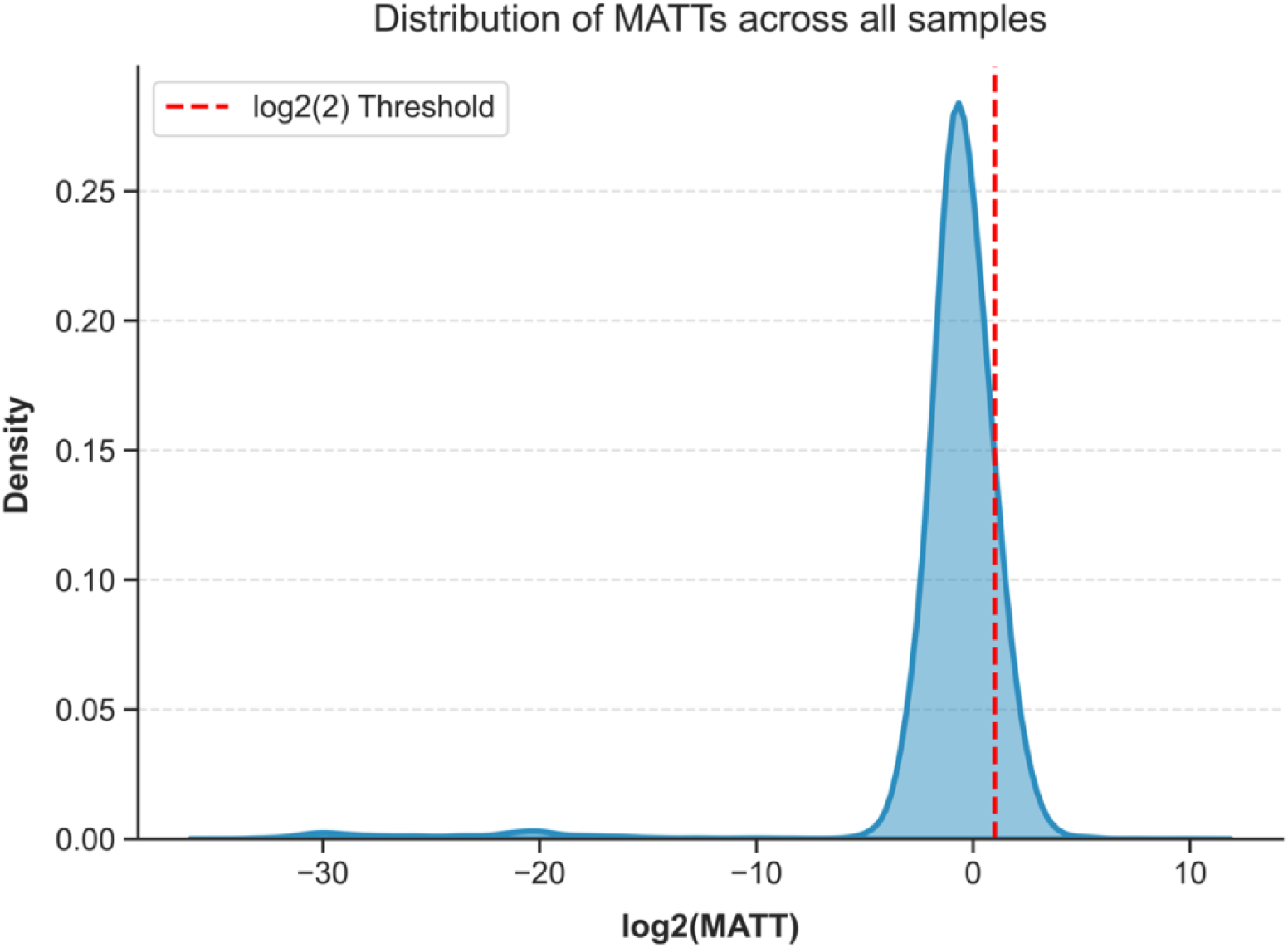
Distribution of MATT threshold values across all Bgee 16 bulk RNA-seq libraries.

**Supplementary figure 2:**
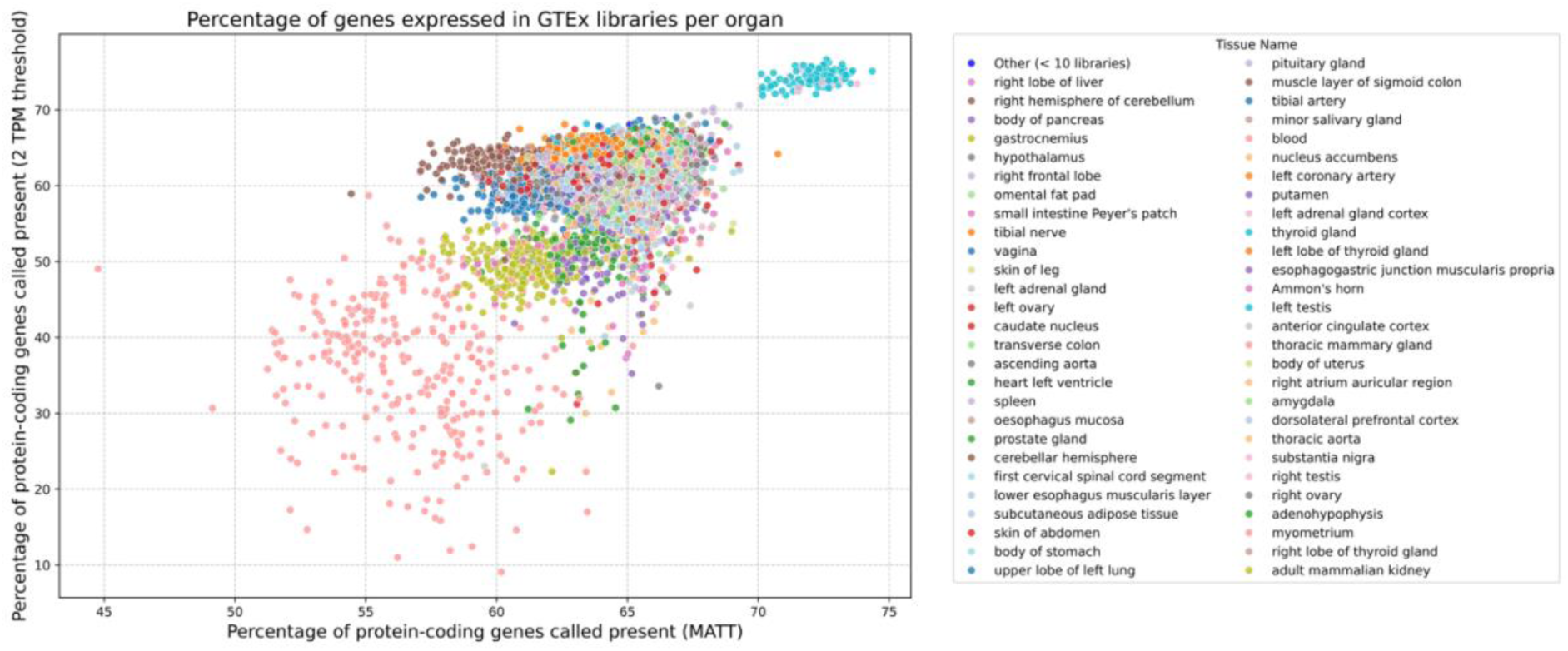
Comparison of proportions of protein-coding genes called expressed using MATT versus a fixed threshold of 2 TPM for all GTEx libraries in Bgee.

**Supplementary figure 3:**
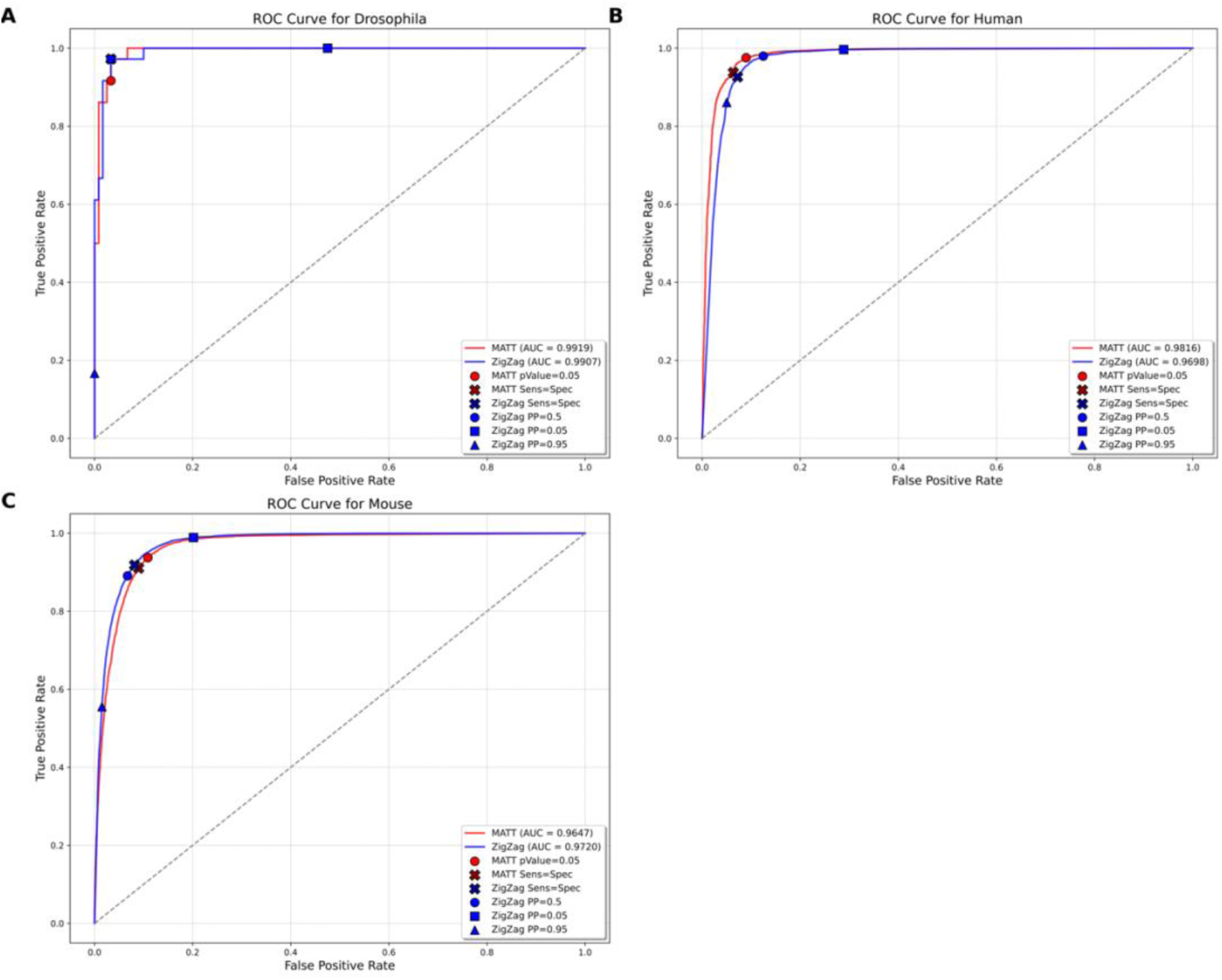
Comparison of the performance of MATT and zigzag to call active expression on a combination of libraries for 3 model species (*Drosophila melanogaster*, Human and Mouse). **(A)** ROC curve of the calls of expression using 9 Drosophila testis libraries. **(B)** ROC curve of the calls of expression for 67 human lung libraries. **(C)** ROC curve of the calls of expression for 330 mouse liver libraries. The red lines show the performance of the MATT-based method and the blue lines show the performance of zigzag. For each method we estimated the inflection point (black cross), and we show the standard thresholds for each method as colored dots.

**Supplementary figure 4:**
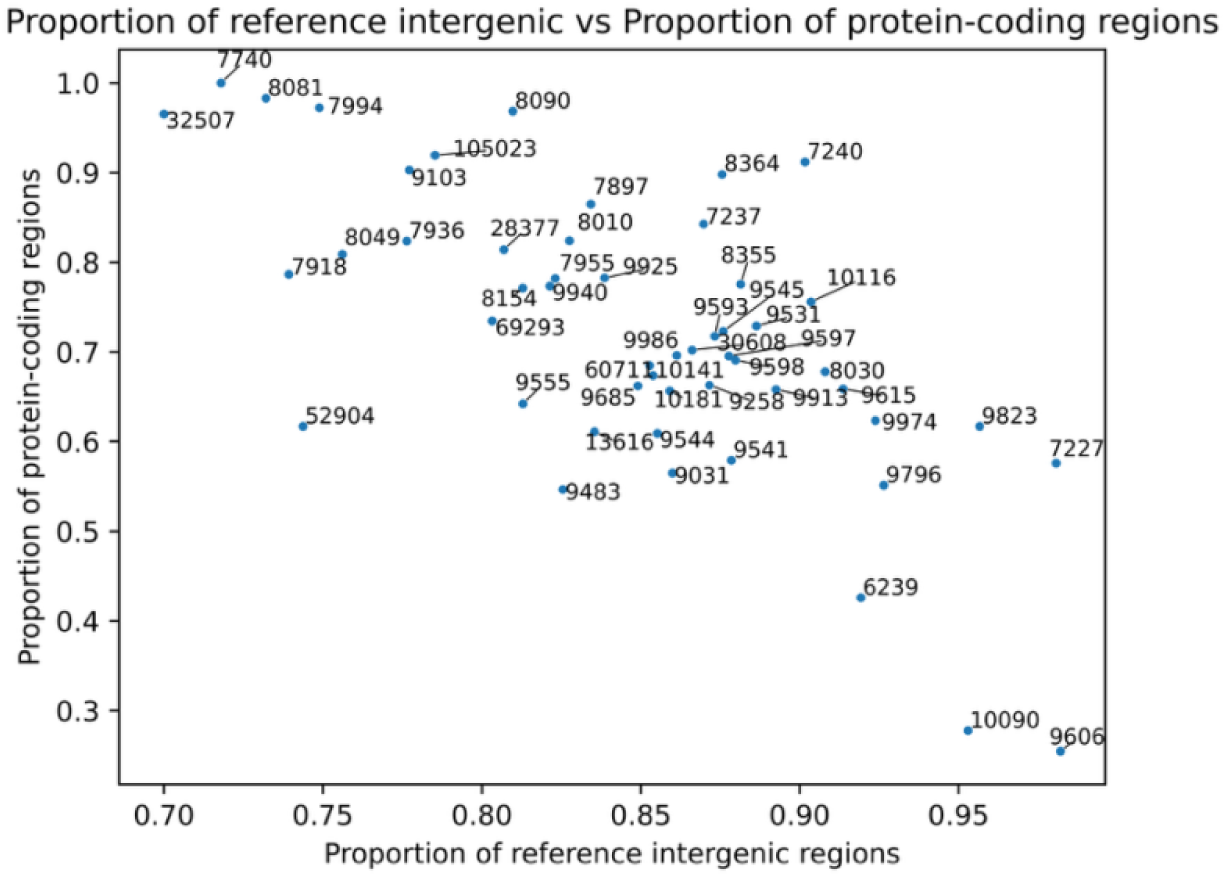
Scatter plot of the proportion of intergenic regions selected to be reference intergenic (among all intergenic regions considered) against the proportion of protein-coding genes among all genes, in each 52 species in Bgee.

## Supplementary Files

Files added to zenodo: https://zenodo.org/records/19347960

## Notes

https://github.com/mrrlab/Robust_data_driven_gene_expression_inference_code

https://zenodo.org/records/19347960

### Competing Interest Statement

The authors have declared no competing interest.

### Summary of Updates

The determination of the reference intergenics was changed to select intergenic regions close to genes. All analyses were revised. The manuscript was tightened.

